# Dual antibacterial properties of copper coated nanotextured stainless steel

**DOI:** 10.1101/2023.10.19.563111

**Authors:** Anuja Tripathi, Jaeyoung Park, Thomas Pho, Julie A. Champion

**Affiliations:** School of Chemical and Biomolecular Engineering, Georgia Institute of Technology, 950 Atlantic Drive, Atlanta, Georgia 30332, United States

## Abstract

Bacterial adhesion to stainless steel, an alloy commonly used in shared settings, numerous medical devices, and food and beverage sectors, can give rise to serious infections, ultimately leading to morbidity, mortality, and significant healthcare expenses. In this study, we have demonstrated Cu-coated nanotextured stainless steel (nSS) fabrication using electrochemical technique and its potential as an antibiotic-free biocidal surface against Gram-positive and negative bacteria. As nanotexture and Cu combine for dual methods of killing, this material should not contribute to drug resistant bacteria as antibiotic use does. Our approach involves applying a Cu coating on nanotextured stainless steel, resulting in a antibacterial activity within 30 minutes. We have performed comprehensive characterization of the surface revealing that the Cu coating consists of metallic Cu and oxidized states (Cu^2+^ and Cu^+^). Cu-coated nSS induces a remarkable reduction of 97% in Gram-negative *Escherichia coli* and 99% Gram-positive *Staphylococcus epidermidis* bacteria. This material has potential to be used to create effective, scalable, and sustainable solutions to prevent bacterial infections caused by surface contamination without contributing to antibiotic resistance.

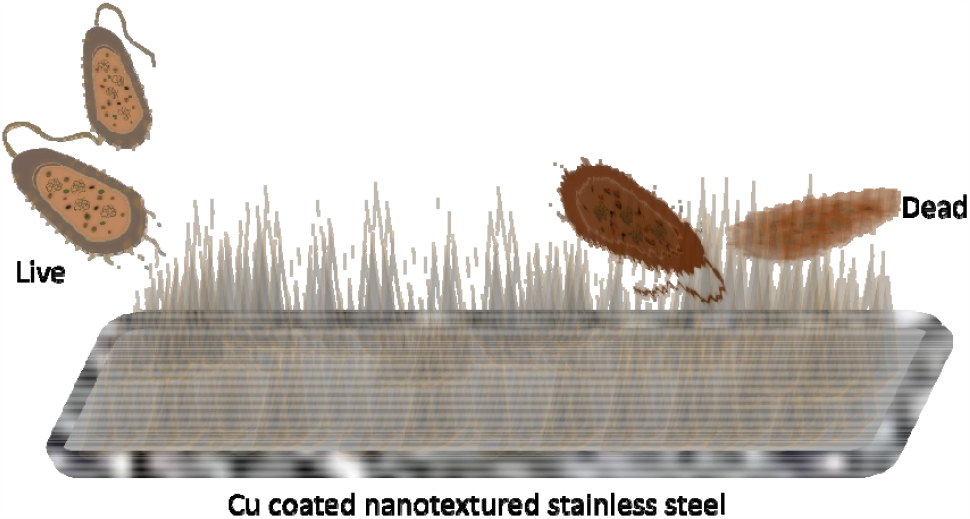

## Introduction

The presence of harmful microorganisms on the surfaces we touch every day, especially in healthcare settings with vulnerable patients, necessitates new approaches to control bacterial contamination and the spread of infection. Modifying surfaces with antibiotics has addressed this concern to a certain extent, but continuous usage of antibacterial agents can lead to development of drug-resistant bacteria[1]. In 2019, drug-resistant bacterial infections caused approximately 1.27 million deaths worldwide[2]. Both Gram-negative and Gram-positive bacteria are health threats, but the outer membrane of Gram-negative bacteria makes them more difficult to kill and more likely to develop resistance[3]. Therefore, there is a significant need to develop antibacterial surfaces effective for both Gram-positive and Gram-negative bacteria. Various approaches have been explored to create bactericidal surfaces, such as coating with antifouling materials, which prevents the adhesion and growth of fouling pathogen substances. However, these coatings often contain toxic biocides such as tributyltin, dichlofluanid, diuron, causing serious health and environmental concerns[4]–[9]. Another approach focuses on developing antibacterial surfaces with inherently antibacterial materials, such as silver and copper, which have antifungal, anti-inflammatory, antiviral, and antimicrobial properties[10], [11]. Previous studies have investigated antibacterial activities of Cu including contact killing, DNA damage, membrane depolarization, and reactive oxygen species generation[12]–[16]. Nevertheless, due to its high cost, copper cannot be extensively employed in bulk. Moreover, using a Cu-coated surface faces the challenge of gradual copper leaching, which diminishes its antibacterial properties over time. Recently, natural antibacterial surfaces, such as microtopographical lotus leaves[17], have inspired polymer, metal, and metalloids nanotextured surfaces[18], [19]. These micro or nanostructures can effectively prevent bacterial adhesion and colonization through direct contact. One such example, reported by Watson et al., used reactive ion etching to create black silicon (bSi) nanowires with diameters ranging from 20 to 80 nm, heights of 500 nm, and inter-wire spacings varying between 200 and 1800 nm. The surfaces were highly bactericidal against both Gram-negative and Gram-positive bacteria, with an average killing rates of up to ∼450,000 cells min^-1^ cm^-2^[20]. Other studies have also explored nanostructured surfaces by employing different cleanroom fabrication techniques such as atomic layer deposition for SiO_2_-ZnO nanowires[21], chemical vapor deposition of amino methyl styrene and vinyl pyrrolidone[22], photo lithography-electron beam lithography and vapor-liquid growth process for SiO_2_ nanowire fabrication[23], and electron beam evaporation for Cr/Zn^0^ thin films[24]. While these materials demonstrated effective antibacterial properties, the feasibility of microfabrication technologies is hindered by complex fabrication processes, lengthy processing times, and high costs. Therefore, it is crucial to identify cost-effective, scalable nanofabrication methods and materials for widespread implementation in combating bacterial infections caused by surface contamination.

Stainless steel 316L (SS316L) is widely used in public settings, including sinks, toilets, surgical tools, and cardiovascular and orthopedic implants, owing to its favorable mechanical strength, corrosion resistance, and biocompatibility[25]. Despite its extensive use, there have been limited studies on the influence of stainless-steel surface topography on bacterial adhesion. Previous investigation revealed that shot peened SS316L surfaces with root mean square (RMS) surface roughness of 29 nm reduced adhesion of Gram-positive bacteria *Staphylococcus aureus* (*S. aureus*) and *Staphylococcus epidermidis* (*S. epidermidis*) but had no significant impact on adhesion of Gram-negative *Escherichia coli* (*E. coli*)[26]. Other surface types with similar roughness parameters showed varying effects on bacterial growth and adhesion. For example, a smooth polymer surface with RMS roughness of 1.4 nm did not have any antibacterial properties. In contrast, a nanotextured polymer surface with an RMS roughness of 13.8 nm effectively inhibited the growth of *S. aureus*, but had less influence on *E. coli* adhesion[27]. These results highlight the significance of surface types, bacterial species, and surface finishing methods in determining bacterial adhesion and growth behavior. In our previous work, we demonstrated the antibacterial properties of nanotextured stainless steel with an RMS roughness of 6.51 nm made by electrochemical etching[28]. This nanotextured stainless steel exhibited reduced *S. aureus* and *E. coli* adhesion and viability while maintaining cytocompatibility with mammalian cells[28].

Nanotextured stainless steel that incorporates inherent antibacterial materials, such as copper, can exhibit dual antibacterial properties. The dual antibacterial activity is due to the small grooves and ridges on the nanotextured surface, which make it difficult for bacteria to attach and spread, and release of copper ions that kill or inhibit the growth of bacteria. In this study, we fabricated nanotextured stainless steel and subsequently coated it with copper using inexpensive, scalable electrochemical techniques for both steps. This approach offers several advantages, including dual-antibacterial properties, affordability, scalability, eco-friendliness, and precise control of surface structures through electrochemical parameters such as potential and current density.

## Experimental Section

### Materials

*E. coli* (BL621) and *S. epidermidis* (PCI 1200) were purchased from ATCC. Luria broth (LB), tryptic soy (TS) media, copper foil, copper nitrate, and nitric acid (ACS reagent, 70%) were purchased from Sigma-Aldrich. Agar and 3,3’,5,5’ tetramethylbenzidine (TMB) were purchased from Fisher-Scientific. SS316L plates (30 × 20 × 0.05 cm^3^) and vinyl insulating tape were purchased from Maudlin Products and 3M, respectively.

### Nanotextured Steel Preparation

Figure 1 below illustrates the electrochemical production of nanotextured stainless steel and its subsequent modification with copper. SS316L was cut into two sizes: 2.5 × 1.5 × 0.05 and 2.5 × 2.5 × 0.05 cm^3^. These served as working and counter electrodes, respectively. The samples were sonicated for 7 min each in acetone, methanol, and isopropanol to eliminate organic contaminants. The samples were rinsed with water to remove organic solvents. A stainless-steel wire was spot-welded onto the SS316L samples to establish electrical connections to the cathode and anode. The working electrode was covered with insulating tape, exposing only an active area of 0.19 cm^2^ for electrochemical surface modification. Diluted nitric acid solution (48 wt.%) was used as an electrolyte (caution! use personal protective equipment and do not mix nitric acid with organics). The working and counter electrodes were separated by 6 cm. Electrochemical surface modification was carried out using a power supply set at 8 V for 30 s. After electrochemical etching, the sample was removed from the electrochemical cell, washed with deionized water, and dried at room temperature.

**Figure 1.**
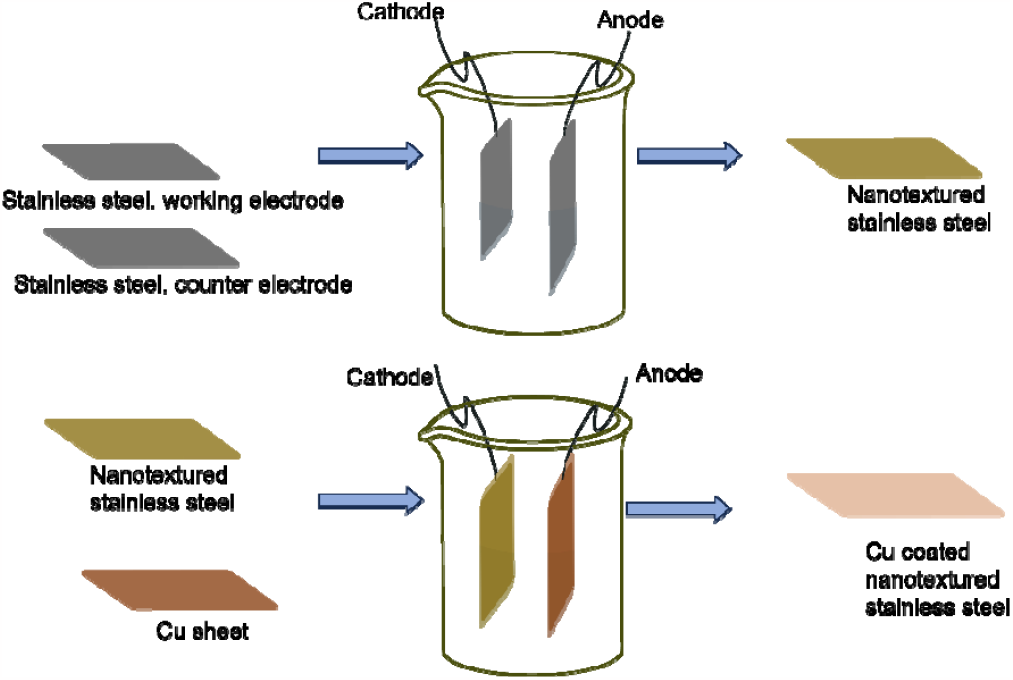
Nanotextured stainless steel fabrication and its modification using Cu by electrochemical techniques.

For the electrochemically deposited Cu on nanotextured stainless steel, we used an electrochemically etched steel as the working electrode and a copper foil as the counter electrode. We applied a voltage of 1 V for different durations of time in 1 M copper nitrate solution. After completing the electrochemical treatment, the coated sample was thoroughly rinsed with deionized water and dried at room temperature.

### Surface Characterization

Surface morphologies of SS316L samples were characterized by scanning electron microscopy (SEM, Hitachi SEM SU8010) at 5 kV acceleration potential, and topographical information was acquired by atomic force microscopy (AFM, Veeco Dimension 3100) using AppNano ACT tapping mode AFM probes (Applied Nanosciences). The surface roughness parameters of stainless steel, nanotextured stainless steel, and copper coated nanotextured stainless steel surfaces were obtained by AFM measurements from scanning a surface area of 4 μm^2^ while avoiding artificial defects. Chemical composition of the sample surfaces was analyzed by X-ray photoelectron spectroscopy (XPS, Thermo Fisher Scientific K-Alpha XPS) with a 400 μm microfocused monochormatic Al Kα X-ray source, which has analysis depth of less than 5 nm. The phase composition of the Cu coated nanotextured steel was determined by X-ray powder diffractometry (Rigaku XRD). The diffraction patterns of samples were recorded in the range of 2□ = 20-80° using a Cu radiation source with fixed power (40 kV, 44 mA). Contact angles of purified water droplets on metal surfaces were monitored using a First Ten Angstrom contact angle goniometer (FTA-200). The Thermo Nicolet 6700 Fourier Transform Infrared Spectrometer (FTIR) was used to measure the infrared energy of Cu.

### Bacterial Cultures and Adhesion Assays

*E. coli* and *S. epidermis* were used as model microorganisms for bacterial assays. All metal samples were sterilized in an autoclave at 15 psi and 121 °C for 20 minutes. The unmodified surface of samples was masked with tape to prevent contamination. These samples were then transferred into 6-well cell culture plates and incubated with 5 mL of bacterial solution. *E. coli* and *S. epidermis* were cultured in LB media and TS media, respectively. The working bacterial solutions had an optical density (OD) of 0.3 at 600 nm unless otherwise mentioned, approximately equivalent to 5 × 10^7^ cells/mL. The samples were cultured with the bacteria for 24 hours in a static incubator at 37 °C in a humidified environment. To quantify the number of adhered *E. coli* and *S. epidermidis* cells on each metal surface, colony-forming units (CFUs) were counted using the spread plate method. After the 24-hour incubation period concluded, the samples underwent five rinses with phosphate-buffered saline (PBS). Following the removal of the tape, they were washed once more before being transferred into a 50 mL tube containing 5 mL of fresh PBS. Each sample was sonicated for 7 minutes and vortexed for 20 seconds to release bacteria from the sample surface into the solution. A series of diluted solutions in PBS was prepared by transferring 25 μL of the resuspended cell solution into 225 μL of fresh PBS, resulting in a 10^−1^ dilution. 25 μL from first dilution was added to 225 uL of nutrient media on a 96-well plate to yield a total volume of 250 µl. The plate was covered with its lid and incubated in a plate reader (Synergy 2, Multi-Mode Microplate Reader, BioTeck) at 30°C. The plate was shaken for 10 s before reading optical density at 600 nm every 5 min over a period of 20 h. Further dilutions (10^−1^ to 10^−9^) were made in PBS in 96-well plates. Then, 30 µL of each diluted solution of *E. coli* and *S. epidermidis* was spread onto LB or TS agar plates, respectively. After 24 hours of incubation at 37 °C, the bacterial colonies on each plate were counted. The number of bacteria (CFU) per sample was calculated by dividing the number of colonies by the dilution factor, multiplying by the amount of cell suspension plated to agar (30 µL). The percentage reduction of bacterial adhesion is calculated by subtracting the number of bacterial colonies on the test surface from the number of bacterial colonies on the control surface, dividing this number by the number of bacterial colonies on the control surface, and multiplying by 100. To visualize bacterial adhesion on the metal surface using scanning electron microscopy (SEM), all metal samples underwent the same procedure described above for incubation with bacteria. After incubation, samples were washed three times with PBS, fixed with a 2.5% glutaraldehyde solution for 1 hour, and dehydrated using a series of ethanol concentrations in distilled water (50%, 70%, 90%, and 100% ethanol for 20 minutes each). The dehydrated samples were dried overnight using hexamethyldisilazane (HMDS) and then coated with gold (approximately 7 nm thickness) using a Quorum Q-150T ES Sputter Coater. The samples were examined using a Hitachi SEM SU8010 at a 5 kV acceleration potential.

### Confocal Laser Scanning Microscopy for DNA Degradation

TUNEL assay was used to detect cell apoptosis using the CF605R TUNEL Assay Apoptosis Detection Kit (Biotium) according to the manufacturer’s instructions. To characterize DNA degradation, steel samples were incubated with bacteria as described above in a static incubator (37 °C, humidified environment) for 30 min. Then, samples were washed with PBS twice, fixed with 4% formaldehyde for 30 min, washed with PBS, then permeabilized with PBS containing 0.2% Triton X-100 for 30 min at room temperature. Samples were finally washed and stained using TUNEL staining buffers while being protected from light. Glycerol mounting media was added onto the stained samples and a #1.5 coverglass was placed (Electron Microscopy Sciences). Finally, samples were imaged using a PerkinElmer UltraView VoX confocal spinning disk system microscope.

### Surface Reactivity Analysis

Steady-state catalysis kinetics assessment was performed on nanotextured steel and copper modified nanotextured steel to catalyze reaction of varying concentrations of TMB under ambient conditions for 15 min. The absorbance values were recorded at 652 nm, and the kinetic catalysis parameters were obtained by linear fitting of the Lineweaver-Burk double-reciprocal plot. The maximal reaction velocity (Vmax) were extracted from:

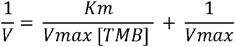

where, V is the initial reaction velocity, and [TMB] refers to the substrate concentration.

### Flow Cytometry for Reactive Oxygen Species Analysis

To assess the effect of metal on the generation of bacterial reactive oxygen species (ROS), bacteria were incubated on metal surfaces as described above for 24 hours at 37°C. Then, the metal surfaces were rinsed with 1 mL of PBS before being scraped to detach and collect *E. coli* and *S. epidermis* in 200 µl of PBS. Detached bacteria were loaded onto a 96-well plate, and the cells were stained with CellROX Deep Red reagent at a final concentration of 250 µM to detect ROS as per manufacturer’s instructions (ThermoFisher Scientific). Following incubation for 45 minutes at 37°C, the cells were analyzed by Cytoflex flow cytometer (Beckman Coulter).

## Results and Discussion

Fabrication and Characterization of Nanotextured Stainless Steel

Nanotextured stainless steel (nSS) samples were prepared by etching stainless steel at 8 V for 10 s to 90 s, and the resulting morphologies are shown in Fig 2 (a, b) and Fig S1. The electrochemical etching time interval was optimized for the desired nanostructure formation, resulting in a distinct morphology of the developed electrochemically treated steel compared to the control sample. We did not observe any noticeable etching within 10 s, but the etching amount significantly increased as the time progressed to 30 s and beyond. Nanopores were distributed regularly over the surface of the steel. For 30 s etching time, the pore size distribution of 20 nm – 30 nm and vertical structure of approximately 30 nm high sharp nano protrusions, which shows a similar morphology as fabricated using clean room techniques[29], [30].

**Figure 2.**
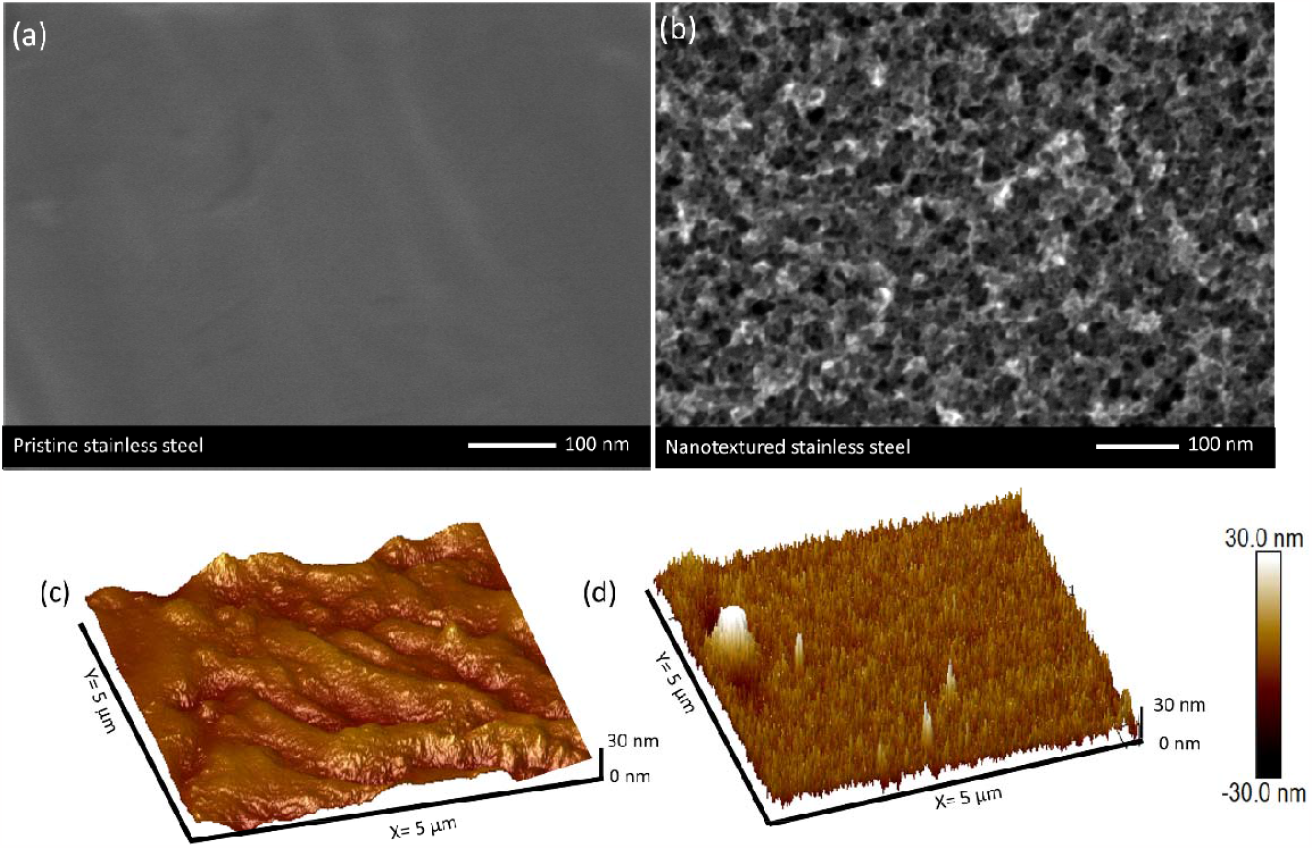
SEM (a, b) and AFM (c, d) images of pristine stainless steel (a, c) and nanotextured stainless steel (b, d) etched for 30 s at 8 V.

Next, we deposited Cu on nSS for various time intervals ranging from 4 min to 15 min at 0.3 A current, as shown in Fig S2. The current causes the positively charged Cu ions to migrate to the cathode (nSS), resulting in different oxidation states of Cu deposited on nSS, as given in the reactions below. Initially, at a coating time of 4 min, SEM images revealed a copper deposited on the textured steel surface. The coating appeared as small islands or particles, indicating the initial nucleation and growth of the coating. The longest deposition time of 15 minutes resulted in a highly uniform and dense layer of copper, completely covering the nSS surface. Hence, we chose two-time intervals for further characterization of Cu deposited nSS, visualized in Fig S3. Cu_4min is a 4-min coating with a noticeable amount of copper islands but majority of exposed nSS surface. Cu_15min is a 15-minute coating with a near continuous, layer on the nSS surface.

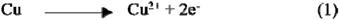

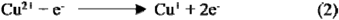

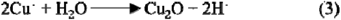

The crystal phase structure of bulk copper was investigated using X-ray diffraction (XRD) in the 20-80° range. Fig S4 shows the XRD pattern, with the highlighted peaks at 2□ = 43.4°, 50.4°, and 74.2°. These peaks correspond to the (111), (200), and (220) crystallographic planes, respectively[31]. Since Cu and Austenite coexist within identical planes, in the case of SS and nSS, these planes might align with Austenite[32], [33]. The increased intensity of the diffraction peaks for Cu_4min and Cu_15min confirms Cu existence, and a larger amount of copper deposition over time. The dominant (111) peak shows that the Cu grains have undergone preferential reorientation in one direction during electrochemical deposition. Furthermore, FTIR spectrum of Cu_4min and Cu_15min in Fig S5 also shows the stretching vibrations of Cu_2_O at 677 cm^-1^, which indicates the Cu/Cu_2_O presence on nSS[34].

The surface chemical composition and elemental valence states of the Cu were investigated by X-ray photoelectron spectroscopy (XPS), and the results are shown in Fig 3. We compared the Cu elemental state between pristine SS and electrochemically treated nSS, Cu_4min, and Cu_15min. Pronounced peaks for Cu^0^ are present in all samples at 932.4 eV and 952.8 eV. SS and nSS showed similar spectra and the low intensity of Cu^0^ present in SS/nSS indicates trace amounts of Cu. The high intensity of Cu^0^ peaks demonstrates the copper coated surface using electrochemistry. The distinct oxidation states were also observed at 934.6 eV and 954.5 eV, corresponding to Cu^2+^ and Cu^+^ respectively, along with their associated satellite peaks at 942.1 eV, 944.2 eV, and 962.9 eV. The charges on Cu were formed during the electrochemical Cu deposition, as given in reactions (1)-(3).

**Figure 3.**
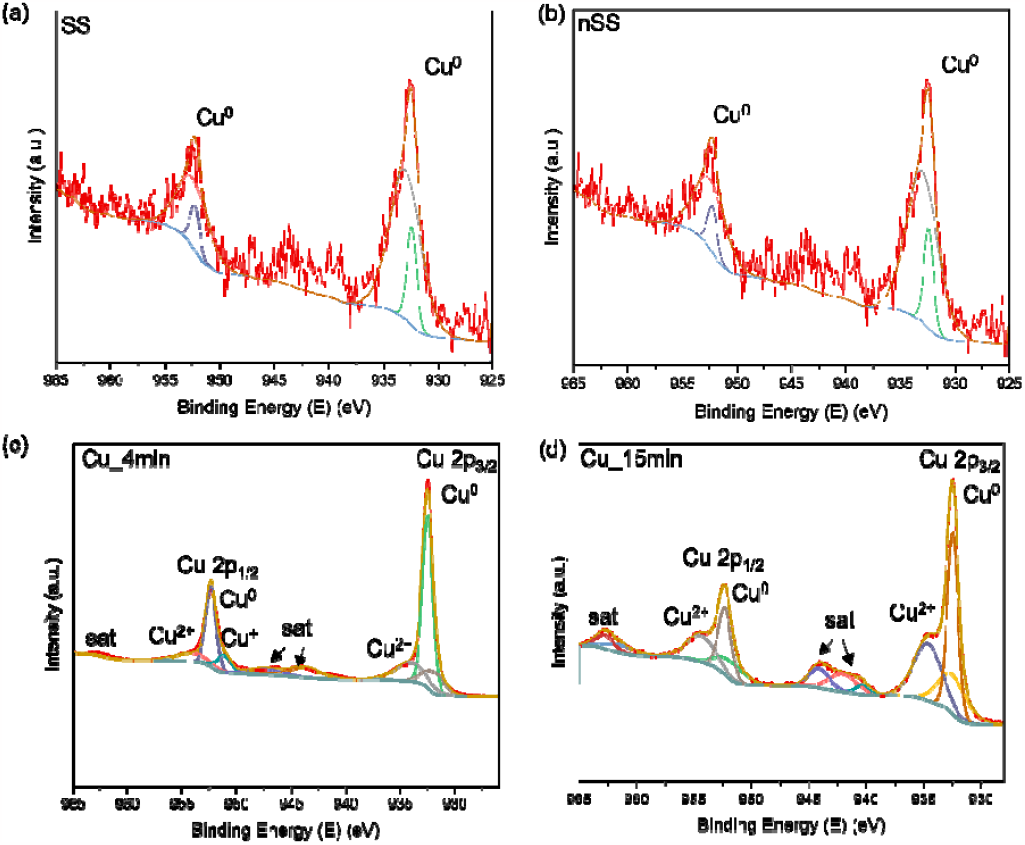
XPS spectra of stainless steel (SS), nSS, and copper coated nSS for 4 minutes and 15 minutes.

To analyze the wetting behavior, contact angle tests were performed on all surfaces (Fig S6). Among them, the hydrophilic stainless-steel surface exhibited the lowest contact angle (60.8° ± 1.21), indicating a strong attraction to water molecules. On the other hand, nSS surfaces demonstrate a much higher contact angle (108.95° ± 0.42) due to increased surface hydrophobicity caused by the nanoprotrusive features. Coating with copper reduced the contact angle somewhat, but the Cu_4min (98.97° ± 1.32) and Cu_15min (96.76° ± 1.84) surfaces still displayed a hydrophobic nature. Hydrophobic surfaces can create an inhospitable environment for bacteria to thrive, as their growth often depends on the presence of moisture[35].

### Generation of Reactive Species

To understand the surface generation of reactive oxygen species (ROS), a subset of free radicals, we conducted kinetic studies utilizing TMB dye. TMB dye can undergo oxidation, transitioning from a colorless state to a blue one in the presence of ROS. Some metal surfaces can catalyze the conversion of dissolved oxygen in water to ROS. These ROS can cross the membranes of bacteria and damage DNA, lipids, and proteins[36]. To measure how well the nSS and Cu-coated nSS surfaces catalyze the production of ROS, we determined the maximum initial velocity of the reaction as a measure of the catalytic activity. Here, the reaction occurs in two steps: 1) The catalytic active sites of metal surface can catalyze ROS generation, 2) ROS oxidize colorless TMB to blue colored TMB, which can be measured at 652 nm. At low substrate concentrations, the reaction rate (V) showed a linear relationship with the substrate concentration [TMB], indicating first-order kinetics (Figure 4). Conversely, at high concentrations, the reaction rate (V_max_) was limited by the availability of active sites on the catalyst, following zero-order kinetics where V is independent of substrate concentration. A catalyst with high effectiveness exhibits a high V_max_, representing a rapid reaction rate, indicating a high number of reactive species generated during the catalytic process. In Figure 4, kinetic studies were performed to compare the catalytic activities of nSS, Cu_4min and Cu_15min. The K_m_ and V_max_ were obtained by fitting the absorbance values for the catalytic reaction products versus time with varying concentrations of TMB in ambient air. While SS did not demonstrate significant catalytic activity (V_max_ = 1.1 nM, Fig S7), nSS (V_max_ = 15 nM/s), Cu_4min (V_max_ = 52 nM/s), and Cu_15min (V_max_ = 55 nM/s) all show strong surface properties for producing reactive species. Overall, this suggests that the nanoprotrusive features of nSS have significantly higher catalytic activity than SS, which was further improved by the presence of Cu on nSS.

**Figure 4.**
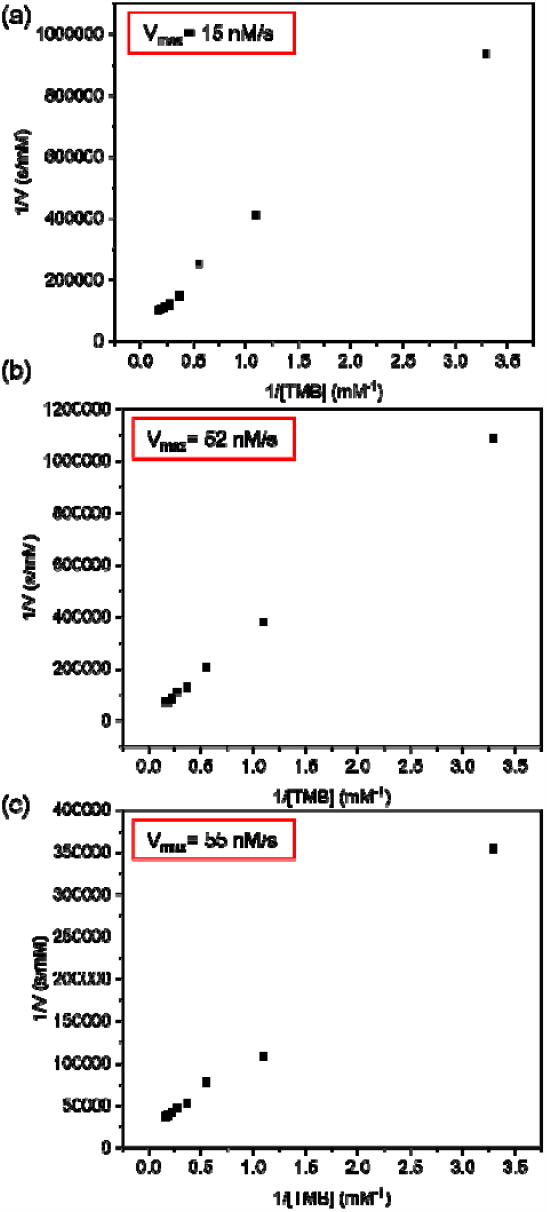
Steady-state kinetic results using Lineweaver-Burk plot for nSS (a), Cu_4min (b), Cu_15min (c), with varying concentrations of TMB at ambient conditions. The reaction time was 15 min and absorbance values were measured at 652 nm.

Next, we measured the Cu leaching in LB media and TS media after 24 h incubation using a Cu calibration for each media, as shown in Fig S8 and S9. As expected, Cu_4min leaches less Cu than Cu_15min for both LB and TS media. This demonstrates that Cu ions are naturally released into media, which corresponds with the occurrence of bacteria death as evident from Fowler et al.’s study that reported Cu can kill bacteria at concentrations of 90 ppm or higher[37].

### Bacterial Inhibition on Metal Surfaces

We conducted a comparative analysis to evaluate the % reduction in bacterial adhesion for nSS, Cu_4min, and Cu_15min and viability of Gram-negative *E. coli* and Gram-positive *S. epidermidis*, with SS as the control. After incubating metal surfaces in bacteria solution for 24 h, adherent bacteria were collected to determine colony formation and growth (Fig 5, Fig S10). The SS surface exhibited high bacterial growth for both Gram-negative and Gram-positive bacteria, indicating its lack of antibacterial properties. On the other hand, nSS, Cu_4min, and Cu_15min surfaces showed either low bacterial growth or no growth at all. The % reduction in bacterial adhesion for nSS was 92.3% for *S. epidermidis* and 84.5% for *E. coli*, which were enhanced when nSS was modified with Cu for different durations. Cu_4min exhibited 94.6% against *E. coli* and 99.2% against *S. epidermidis*. Similarly, Cu_15min displayed further enhanced % reduction, achieving 96.8% against *E. coli* and 99.6% against *S. epidermidis*. The notable improvements in dual-antibacterial surfaces can be attributed to the combined properties of nanoscale roughness and Cu release. Nanotextures have the capacity to physically harm bacterial cells by either causing physical damage or penetrating the cell wall. Additionally, they can generate ROS, which are highly reactive molecules capable of inflicting damage to bacterial cells. Cu toxicity has been attributed to ROS generation and membrane depolarization, which can induce bacterial stress and disrupt outer membranes[38].

**Figure 5.**
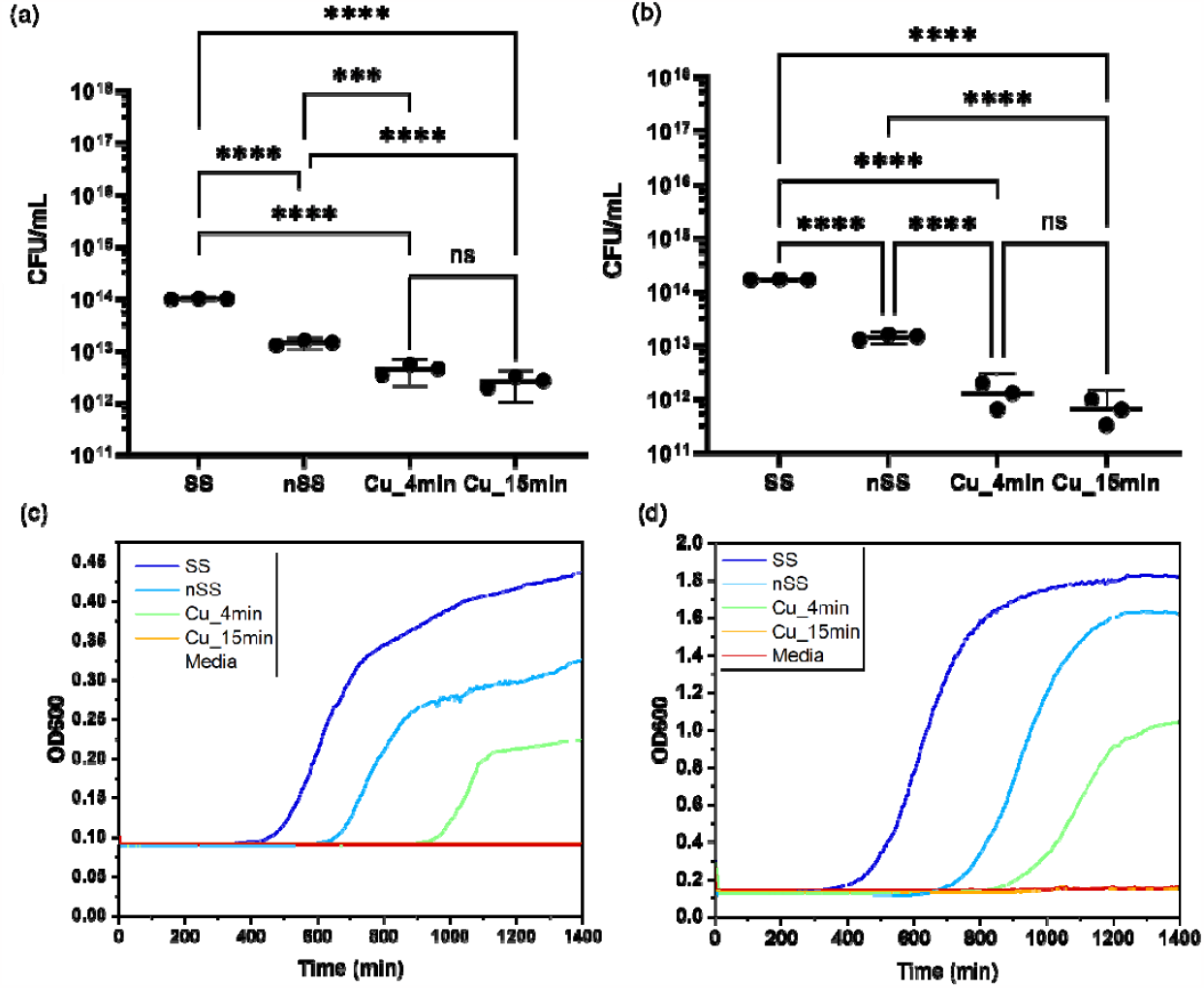
Bacterial growth curves of *E. coli* in LB media (a) and *S. epidermis* in TS media (b). % reduction of bacterial adhesion after incubation for 24 h with nSS, Cu_4min, and Cu_15min (c).

The bacterial membrane plays a crucial role in generating a proton gradient for adenine triphosphate synthesis, the primary energy carrier of cells. The disruption of the membrane interface can result in reduced energy production and physical damage to the bacterial cells [39]. This can be observed in the negative effects on bacterial growth patterns following removal from the nanotexture and copper (Figure 5c,d). For SS, *E. coli* and *S. epidermidis* both exhibited growth after a lag time. However, after exposure to nSS, the growth of both *E. coli* and *S. epidermidis* was inhibited, resulting in increased lag times, and lower OD600 at the end of growth. Exposure to Cu_4min further inhibited growth for *E. coli* and *S. epidermidis* and no growth was observed for exposure to Cu_15min. Altogether, this data indicates that both nanotexture and copper can inhibit bacterial growth by both delaying the onset of exponential growth and reducing the growth rate. These growth curve patterns align with previously reported literature on antibacterial activities of Rh nanoplates[40]. This is consistent with the findings of Hasan et al., who reported that decreasing bacterial growth with increasing Cu and no bacterial growth was observed with copper nanoparticles at a concentration of 60 µg Cu/mL or higher[41]. Moreover, SEM imaging was performed to examine *E. coli* adhesion on SS, nSS, Cu_4min, and Cu_15min, as shown in Figure S11. No morphological alterations were observed in the adhered bacteria on SS, suggesting that there was no bacterial damage. Conversely, wrinkled morphology and a reduced number of bacteria were observed on nSS.

### Assessment of ROS Production and DNA Degradation in Bacteria

As stated earlier, ROS generation from nSS and copper coated nSS surfaces were detected and could be a cause of the decrease in bacteria adhesion and growth. To quantitatively determine the intracellular generation of ROS induced by nSS and Cu-coated nSS, we measured the CellROX Deep Red fluorescence intensity using a flow cytometer. We included untreated bacteria and bacteria treated with peroxide, SS, and copper foil as controls. Both *E. coli* and *S. epidermidis* exhibited high ROS production on nSS surfaces, but not on SS or either copper coated surface (Fig 6a,b). While *E. coli* had a ROS negative population and ROS positive population that was lower than the peroxide positive control, *S. epidermidis* cells all produced more ROS than peroxide treated cells (Fig S12). This is consistent with greater reduction in *S. epidermidis* CFU by nSS compared to *E. coli*. These results indicate that the sharp features of nSS promoted induction of oxidative stress in bacteria, resulting in ROS production, which could cause the reduced CFU and growth observed [42]. The mechanism of antibacterial activity mediated by Cu remains incompletely defined due to conflicting reports on whether Cu ions induce bacterial generation of ROS. Despite evidence of free radical production by all modified surfaces (Fig 4), we did not observe any ROS generation in bacteria cultured on Cu foil, Cu_4min, and Cu_15min (Fig 6a,b, S12). This suggests that the antibacterial activity of copper-modified nSS is not just ROS dependent, and may be mostly promoted by membrane depolarization[43]–[45]. In Fig 3, Cu^2+^ and Cu^+^ was detected by XPS, implying that the positively charged ions may bind to negatively charged domains on the bacterial cell membrane, both outer and inner, thereby reducing the electrical potential difference across the membrane, which could rupture the membrane [46]. It is possible that the process of membrane depolarization occurs more rapidly than the nSS induced cellular production of ROS, which would require physical interaction of the bacteria with the surface, so ROS is only produced in the presence of nSS surfaces without copper. To validate this, we measured the change in membrane potential, which is essential for energy transduction during respiration, after incubating in Cu-modified nSS. We used a BacLight membrane potential kit, which contains DiOC2(3) (3,3’-diethyoxacarbocyanineiodide), a dye that initially emits green fluorescence in polarized cells. However, this green fluorescence decreases upon loss of polarization. It is important to note that the dye is not as accurate in measuring membrane potential in Gram-negative bacteria as it is Gram-positive bacteria[47], which is why this experiment was performed only on *S. epidermidis*. In Figure S13, we observed a decrease in green fluorescence for Cu_15min modified nSS. This shift suggests that the interaction between bacteria and metal allows Cu ions to penetrate the cells, potentially facilitating access to cellular components. Subsequently, intercellular Cu ions may cause irreversible damage and cell death[48]. Figure 6c shows that Cu_15 min exhibited a significant population of cells with membrane depolarization in comparison to Cu_4min, likely due to the higher Cu concentration. Additionally, Cu contact killing might have damaged the outer and/or bacterial membrane, led to accumulation of Cu in the cells, and degraded bacterial DNA [49], [50]. Also, accumulation of intracellular copper ions can scavenge ROS, reducing our ability to measure it [51]. While more studies will be needed to clarify the mechanism underlying antibacterial activity, these results demonstrate that different mechanisms are involved for antibacterial activity via nSS and Cu coated nSS surfaces.

**Figure 6.**
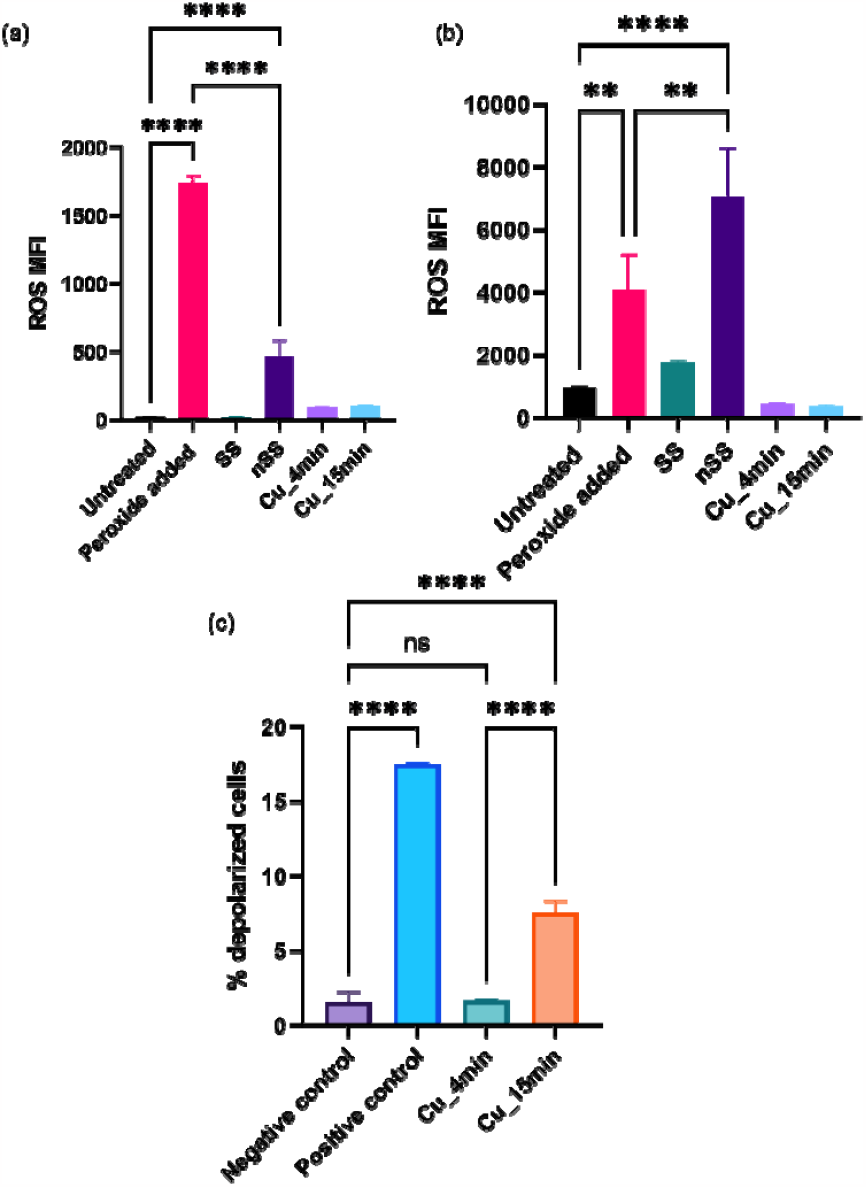
Mean fluorescence intensity (MFI) of *E. coli* (a) and *S. epidermidis* (b) labeled for ROS after incubating with nSS, Cu_4min, and Cu_15min. Untreated cells, peroxide added cells, and SS incubated were taken as control samples. (c) % depolarized *S. epidermidis* cells with Cu_4min and Cu_15min. Untreated cells and treatment with carbonyl cyanide 3-chlorophenylhydrazone (CCCP, a membrane depolarizing agent) were used as negative and positive control samples, respectively. Data represented here as mean ± s.d. **p≤ 0.01 and **** p≤ 0.0001.

Ion contact with bacteria has been linked to mechanisms associated with membrane damage and DNA degradation[52]. Copper is a naturally occurring element that is constantly oxidizing or losing electrons. This process produces copper ions, which are highly reactive and can damage the cell walls of bacteria. When bacteria encounter copper ions, they absorb them and the copper ions can damage DNA, ultimately causing cell death [53]. To investigate the impact of Cu coated nSS on DNA damage, we used the TUNEL (terminal deoxynucleotidyl transferase dUTP nick end labeling) Assay to evaluate DNA fragmentation of bacteria adhered to surfaces (Fig 7). The assay fluorescently labels the 3′-ends of DNA strand breaks, which is a hallmark of apoptosis[54], [55]. SS showed no fluorescence, while nSS shown to have an increase fluorescent signal for DNA fragmentation. nSS was shown to have an increase in both abiotic and intracellular ROS based on the TMB and CellRox assays, respectively, which could contribute to increase DNA fragmentation compared to SS surfaces for both *E. coli* and *S. Epidermidis*. Cu_4min and Cu_15min showed more fluorescence than SS and nSS, indicating that they caused more DNA fragmentation, possibly due to Cu ion interaction with the bacterial cell wall.

**Figure 7.**
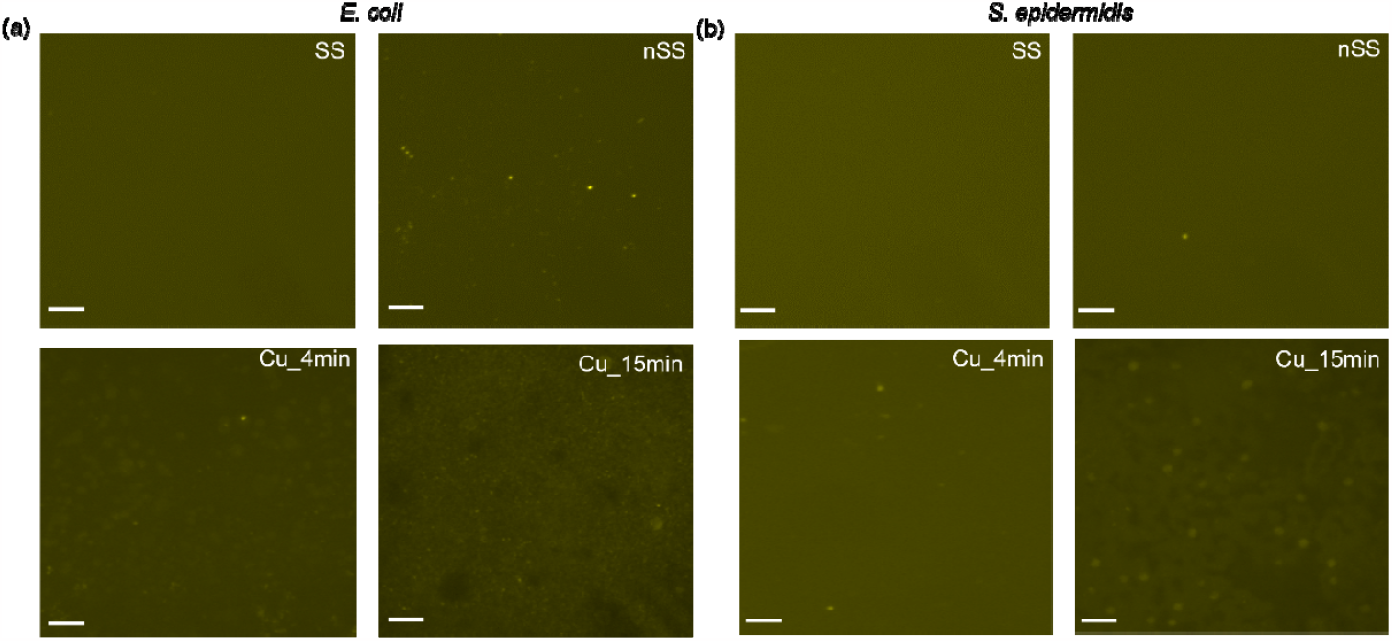
Representative fluorescent micrographs of DNA fragmentation of *E. coli* (a) and *S. epidermidis* (b) cultured for 30 min on SS, nSS, Cu_4min, and Cu_15min and fixed. All scale bars are 30 µm.

## Conclusion

We have demonstrated dual activity surfaces for killing bacteria using copper-coated nanotextured stainless steel. We achieved this by employing an inexpensive electrochemical technique to first create nanotextured surfaces on SS316L, featuring nanopores and nanoprotrusions measuring 20-30 nm, then deposit copper onto the nanotextured surface, resulting in globular or thin film morphology. Both the copper-coated nanotextured stainless steel and the bare nanotextured stainless steel displayed enhanced antibacterial qualities compared to regular stainless-steel surfaces. While the copper will gradually leach out from the metal over time, the underlying nanotextured stainless steel structure will remain intact, ensuring that the surface retains antibacterial attributes even after the copper has been released. This advancement has significant potential for practical usage, as it offers a method to prevent bacterial adhesion and surface contamination without antibiotics. Further, the dual function may not contribute to development of drug resistant bacteria like antibiotics do. The cost-effectiveness and scalability of this surface modification approach makes it practically relevant for larger scale surfaces in public or healthcare settings.

## Acknowledgements

This study was financially supported by Anuja Tripathi’s Presidential Postdoctoral Fellowship at Georgia Institute of Technology. This work was performed in part at the Georgia Tech Institute for Electronics and Nanotechnology, a member of the National Nanotechnology Coordinated Infrastructure, which is supported by the National Science Foundation (Grant No. ECCS-2025462). This work also used core facilities of the Petit Institute of Bioengineering and Biosciences at Georgia Tech. We are thankful to Bradley Park for his kind help with cutting steel pieces for this study.

