## Supplemental file for "Dual antibacterial properties of copper coated nanotextured stainless steel"

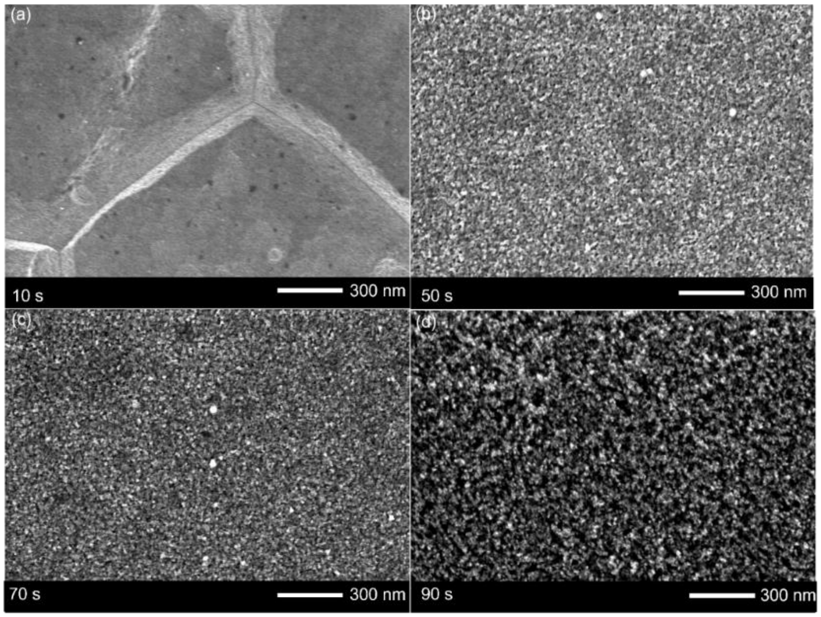


Figure S1: SEM images of electrochemically treated stainless steel at (a) 10 s, (b) 50 s, (c) 70 s, (d) 90 s.

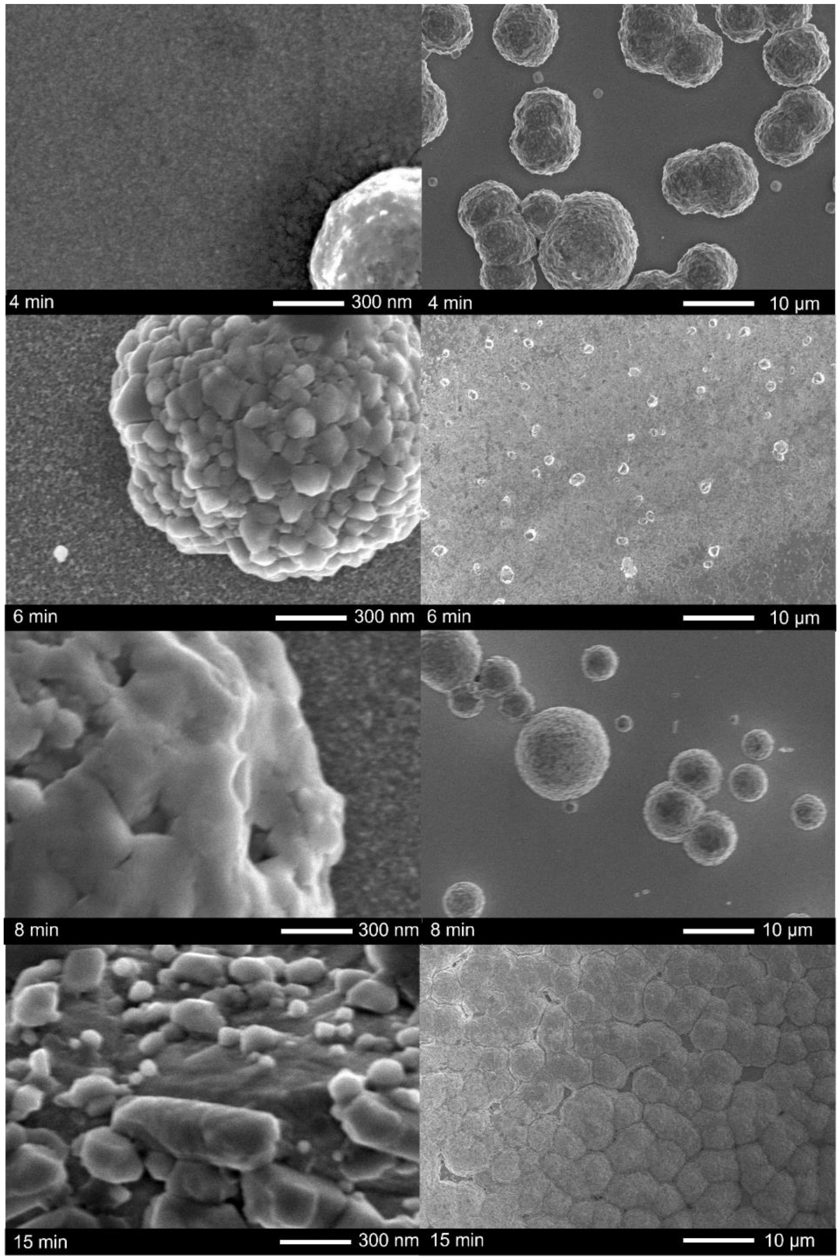


Figure S2: SEM images of copper coated nanotextured stainless steel at different time intervals.

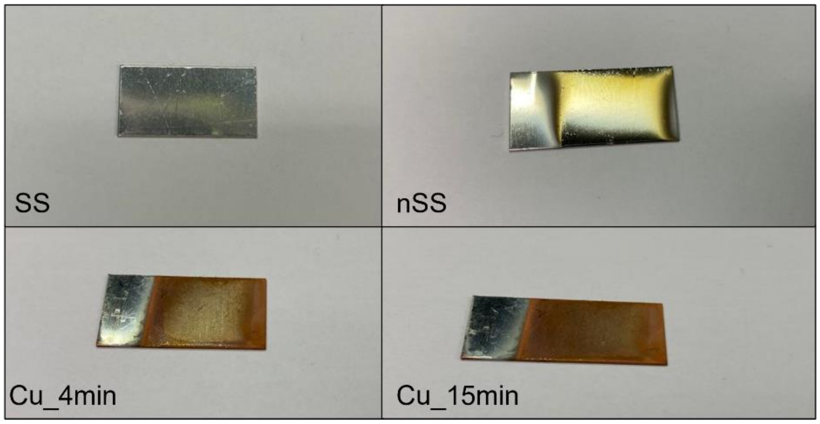


Figure S3: Visual photographs of SS (pristine), nSS, Cu_4min, and Cu_15 min.

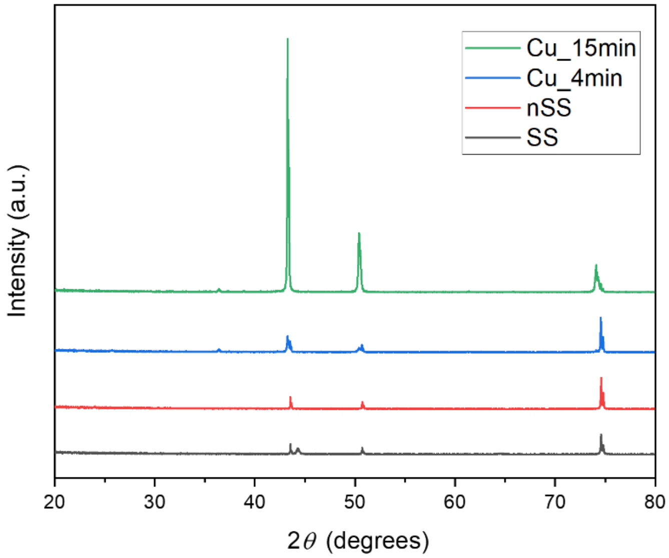


Figure S4: XRD patterns of SS, nSS, Cu_4min, and Cu_15min.

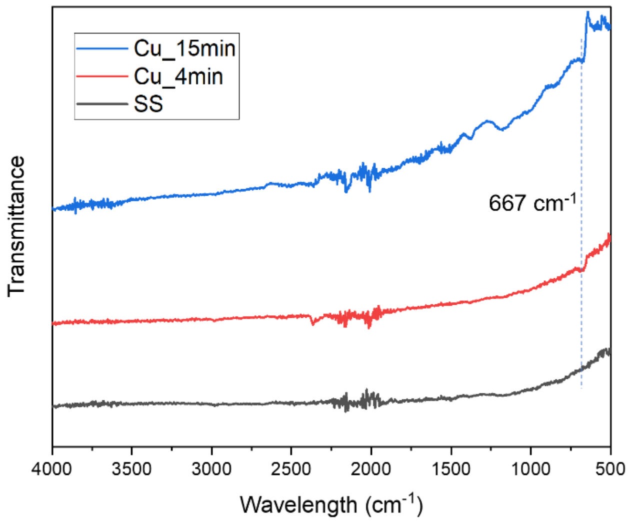


Figure S5: FTIR spectroscopy for SS, Cu_4min, and Cu_15min.

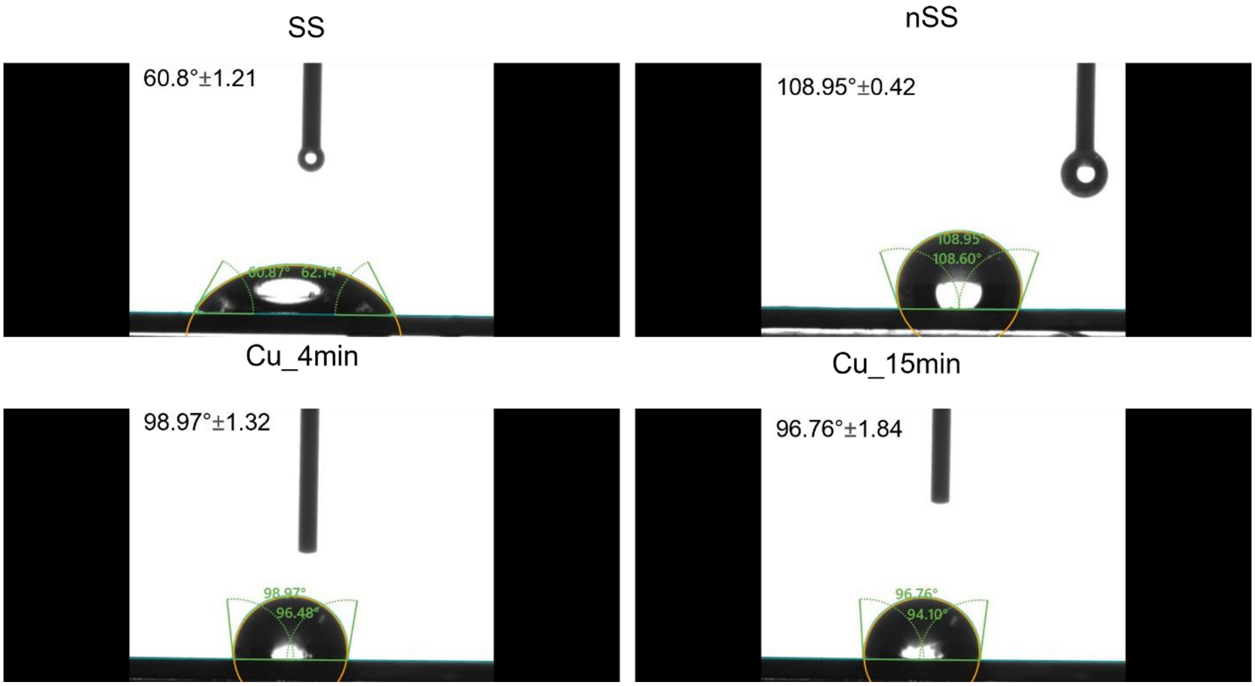


Figure S6: Representative contact angle measurements and average contact angles for SS, nSS, Cu_4min, and Cu_15min.

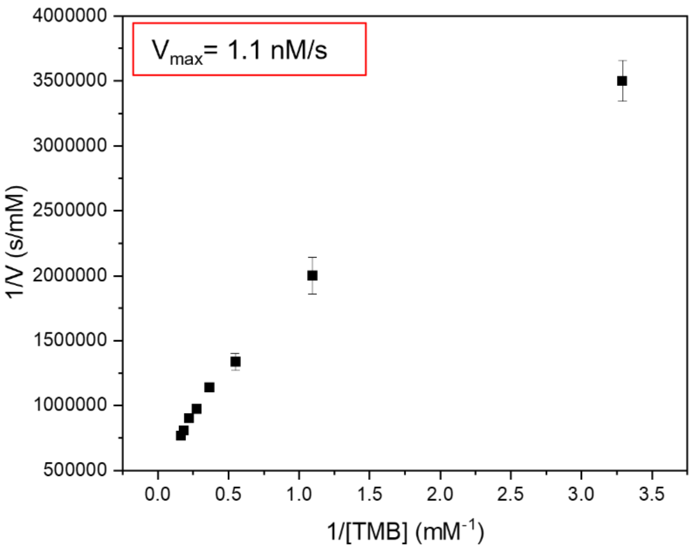


Figure S7: Steady-state kinetic result using Lineweaver-Burk plot for SS with varying concentrations of TMB at ambient conditions. The reaction time was 15 min and absorbance values were measured at 652 nm.

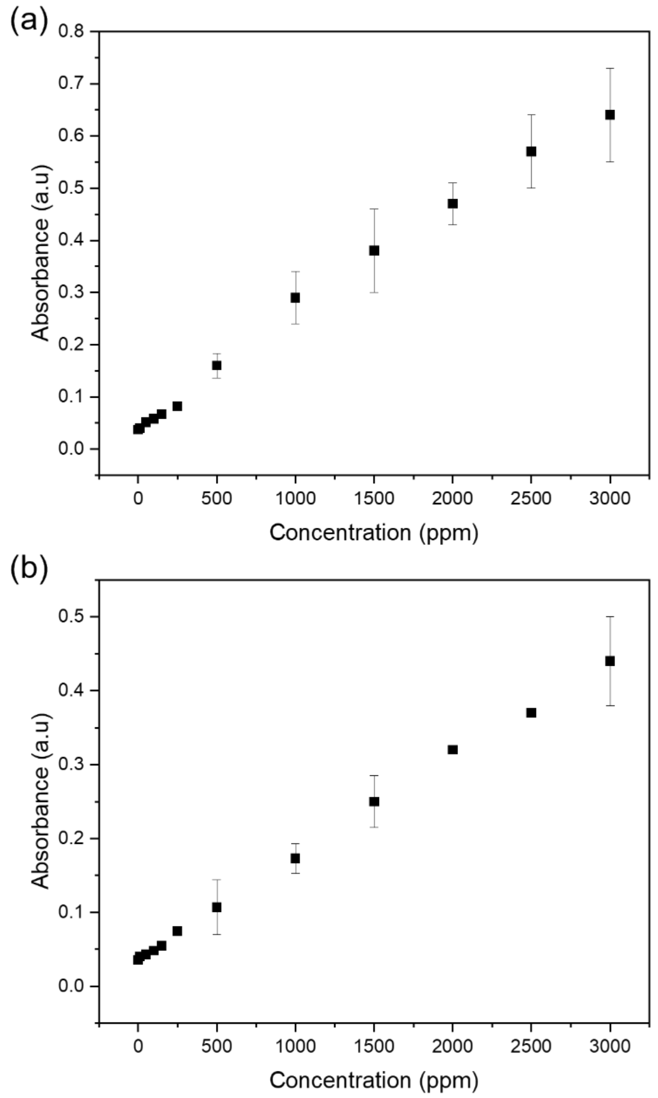


Figure S8: Cu calibration curve in TS media (a) and LB media (b). The calibration equations for Cu in LB media and TS media are [Cu] = 0.0001 Absorbance + 0.0378, $R^2$ = 0.998, and [Cu] = 0.0002 Absorbance + 0.0447, $R^2$ = 0.993, respectively.

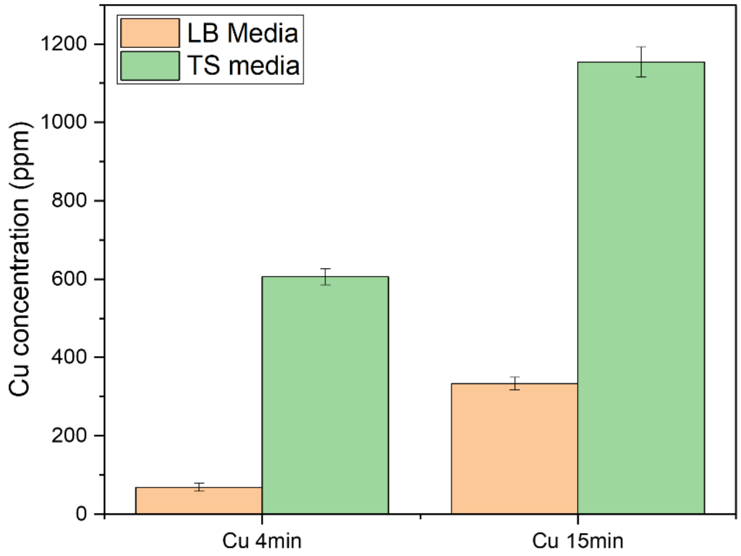


Figure S9: Leached Cu calibration curve for Cu_4min and Cu_15min in TS and LB media after 24 hours incubation at 37°C.

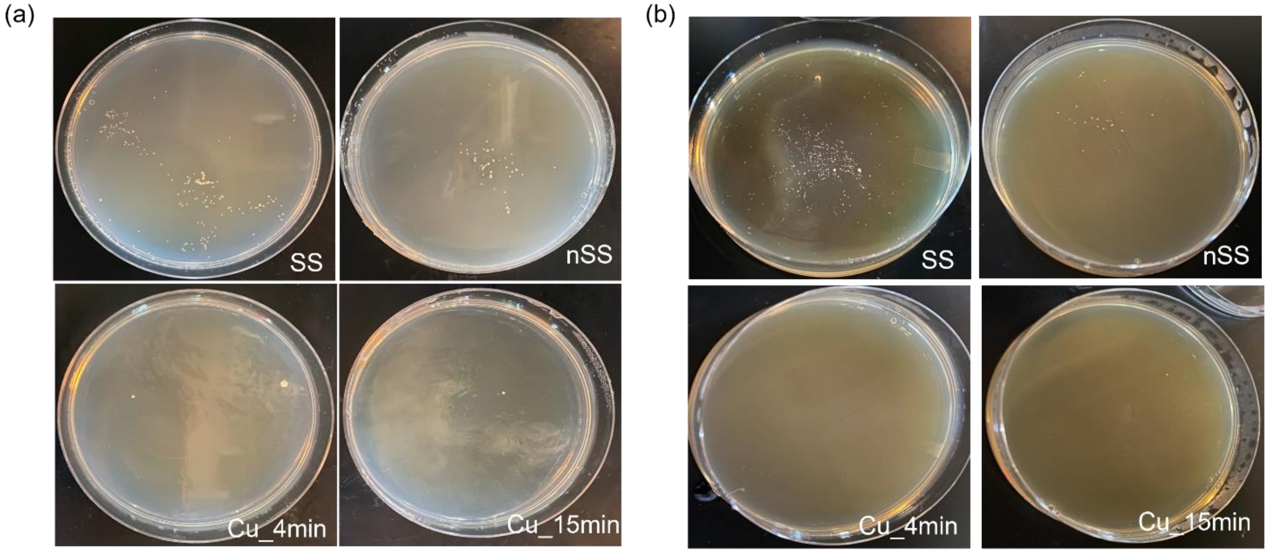


Figure S10: Colony forming units for SS, nSS, Cu_4min, and Cu_15min after incubation for 24 h in *E. coli* (a,c) and *S. epidermidis* (b,d) in $10^{-9}$ dilution

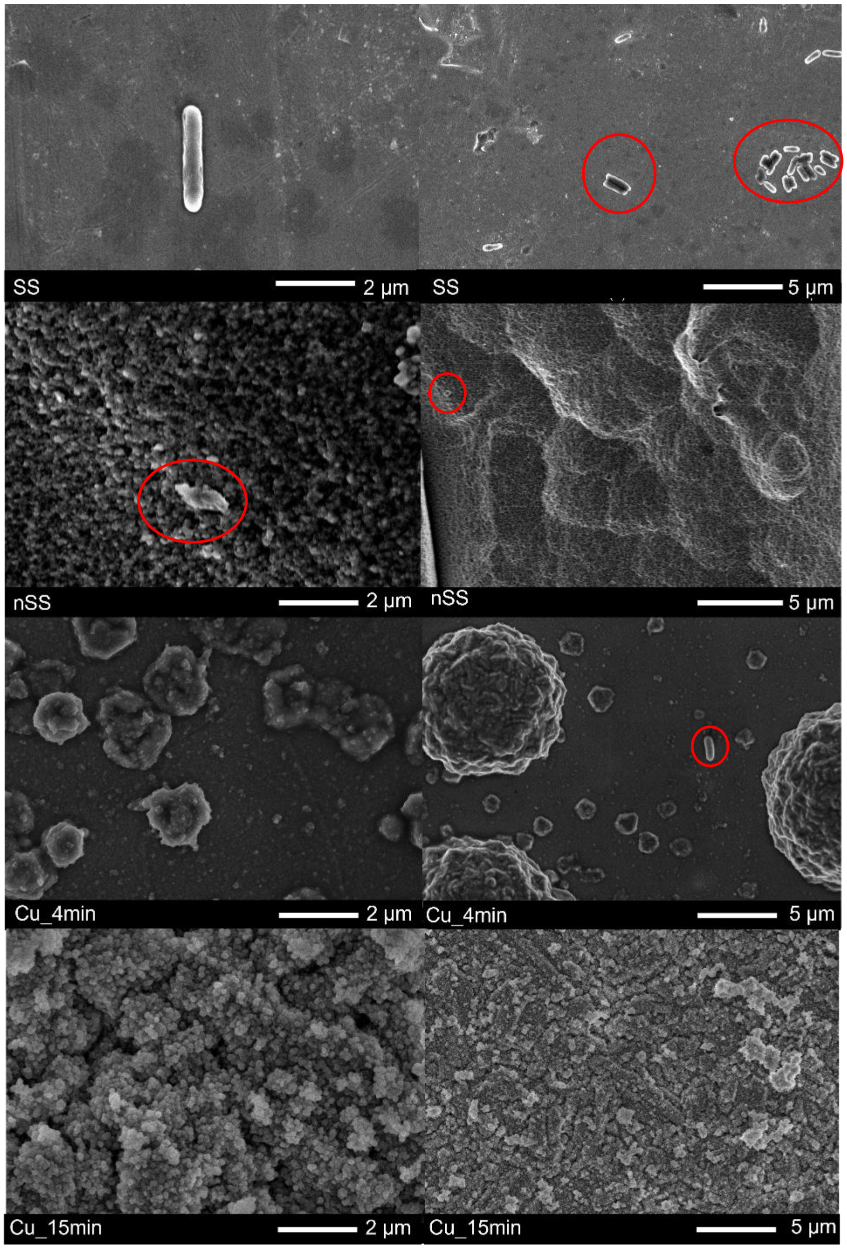


Figure S11: *E. coli* adhesion after incubating on SS, nSS, C_4min, and Cu_15min, for 24 h.

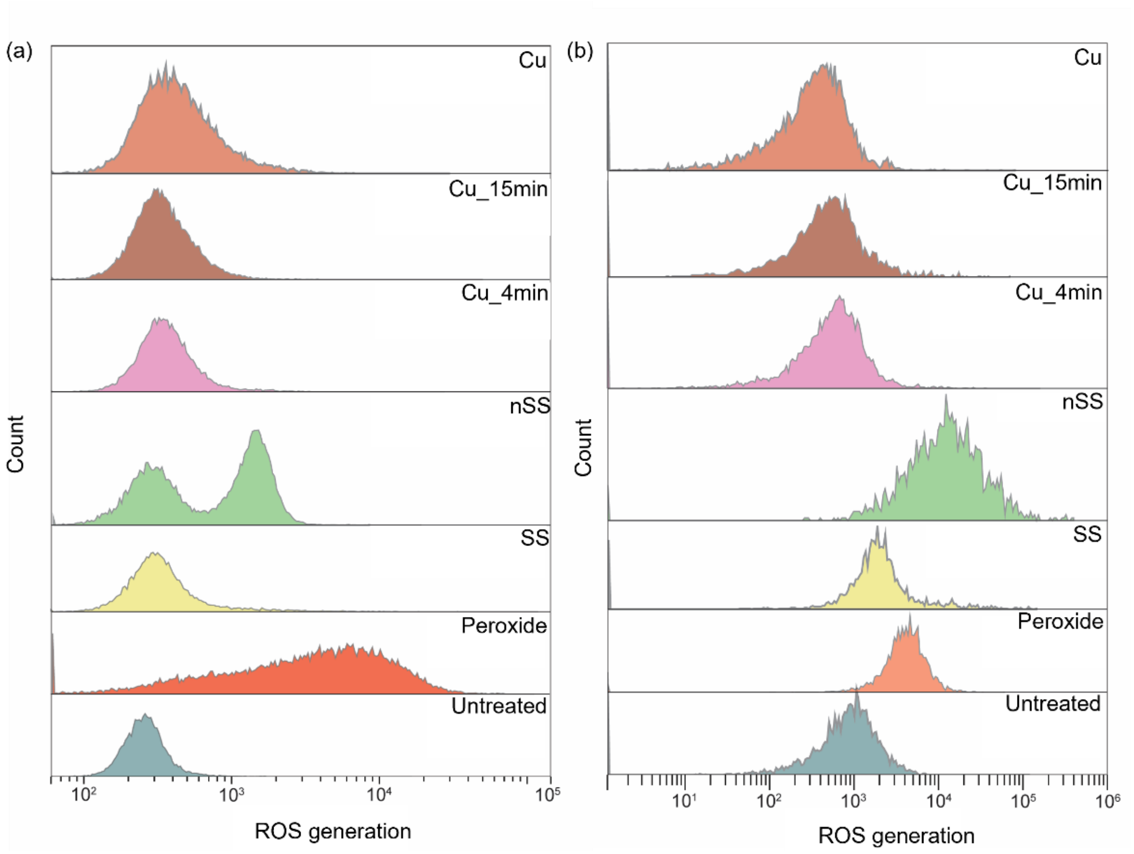


Figure S12: Flow cytometer histograms for ROS fluorescent labelling of *E. coli* (a) and *S. epidermidis* (b) after incubating with nSS, Cu_4min, and Cu_15min. Untreated cells and treatment with peroxide, SS, and pure Cu foil were used as control samples.

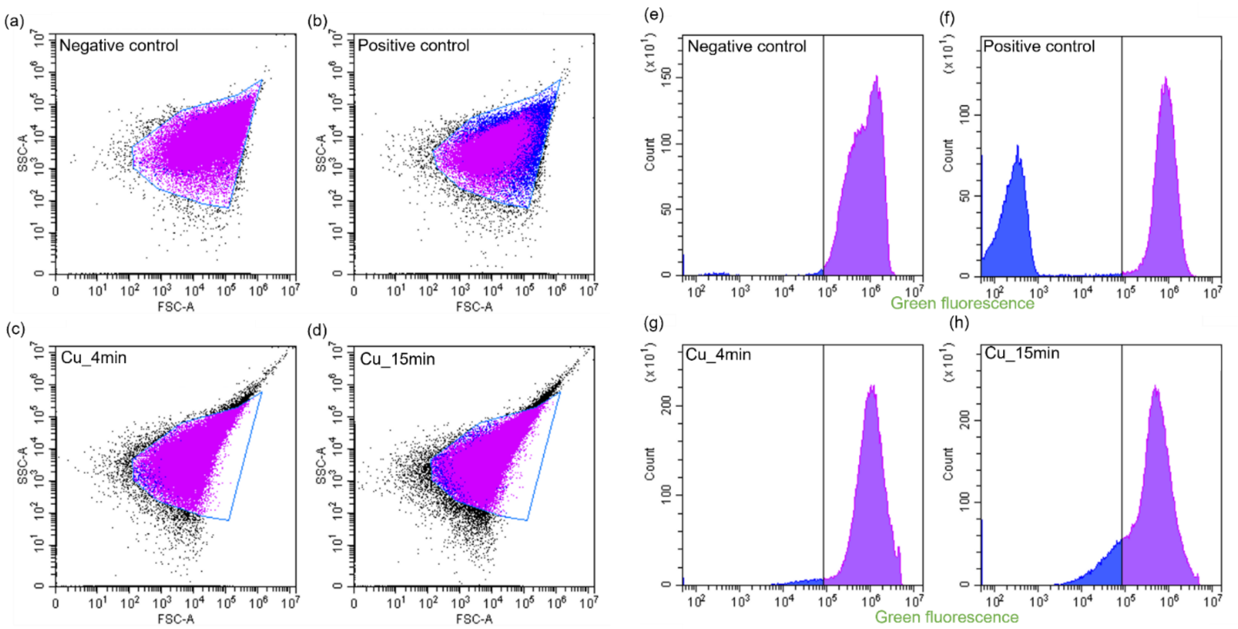


Figure S13: Representative raw data for membrane potential measurement. (a-d) Forward scatter (cell size) vs side scatter (cell granularity) plots for *S. epidermis* cells. Gate for cells was drawn from negative control. Cells exposed to nSS with copper exhibited a shift in these properties. (e-h) Flow cytometer histograms for green fluorescence shift of *S. epidermidis* after incubating with nSS Cu_4min and Cu_15min. Cells positive for membrane depolarization (reported in Figure 6c) are those to the left of the gate (blue population), drawn based on the negative control (majority of cells remain polarized, purple population). Untreated cells and treatment with carbonyl cyanide 3-chlorophenylhydrazone (CCCP, a membrane depolarizing agent) were used as negative and positive control samples, respectively.
